# Functional analysis of V2 protein of Beet curly top Iran virus

**DOI:** 10.1101/2022.09.19.508497

**Authors:** Atiyeh Bahari, Araceli Castillo Garriga, Naser Safaie, Eduardo Rodriguez Bejarano, Ana Isabel Perez Luna, Masoud Shams-Bakhsh

## Abstract

The geminivirus beet curly top Iran virus (BCTIV) is one of the main causal agents of the beet curly top disease in Iran and the newly established *Becurtovirus* genus type species. Although the biological features of known becurtoviruses are similar to those of curtoviruses, they only share a limited sequence identity, and no information is available on the function of their viral genes. In this work, we demonstrate that BCTIV V2, as the curtoviral V2, is also a strong local silencing suppressor in *Nicotiana benthamiana* and can delay the systemic silencing spreading, although it cannot block the cell-to-cell movement of the silencing signal to adjacent cells. BCTIV V2 shows the same subcellular localization as curtoviral V2, being detected in the nucleus and perinuclear region, and its ectopic expression from a PVX-derived vector also causes the induction of necrotic lesions in *N. benthamiana* like the ones produced during the HR, both at local and systemic levels. The results from the infection of *N. benthamiana* with a V2 BCTIV mutant showed that V2 is required for systemic infection but not for viral replication in a local infection. Considering all these results, we can conclude that BCTIV V2 is a functional homologue of curtoviral V2 and plays a crucial role in viral pathogenicity and systemic movement.

## 1. Introduction

Geminiviruses are a group of insect-transmitted plant viruses that cause destructive diseases in major crops worldwide (García-Arenal & Zerbini, 2019; Rojas et al., 2001). The virions of this family are twinned and contain a single copy of circular single-stranded DNA, ranging in size from 2.5 to 3.0 kb. The family *Geminiviridae* includes more than 450 species organized into 14 genera according to the genome organization (one-molecule genomes in monopartite and two-molecule genomes in bipartite) and other biological features, such as host range and type of insect vector (Roumagnac et al., 2022).

RNA silencing is one of the most efficient antiviral defence in plants. It is activated by a viral double-stranded RNA (dsRNA) that is then processed by Dicer-like (DCL) ribonucleases producing 21 to 24-nt primary viral small interfering RNAs (vsiRNAs). Subsequently, in the amplification step, host-encoded RNA-dependent RNA polymerases (RDRs) synthesize dsRNA from viral single-stranded RNA producing secondary vsiRNAs that, together with the primary vsiRNAs are responsible for the spreading of the silencing through the plant. vsiRNAs bind to different Argonaute (AGO)-containing effector complexes and provide targeting specificity for RNA dicing (PTGS or post-transcriptional gene silencing) or chromatin modification (TGS or transcriptional gene silencing). Viruses counterattack by producing one or several proteins that suppress this defence by targeting different steps in the pathway (Bologna & Voinnet, 2014; Borges & Martienssen, 2015; Csorba et al., 2015; Pumplin & Voinnet, 2013).

Geminiviruses must confront both types of silencing, PTGS, and TGS. As counter-defence, they encode more than one viral RNA silencing suppressor, being V2 the proteins exhibiting the stronger PTGS suppression activity (Hanley-Bowdoin et al., 2013; Pooggin, 2013)

V2 (AV2) is a multifunctional protein that has been described as (i) PTGS or a TGS suppressor in most of the geminivirus species tested to date (Amin et al., 2011; Li et al., 2021; Luna et al., 2017; Mubin et al., 2010; Sharma & Ikegami, 2010; B. Wang et al., 2014; Yang et al., 2018; Zhai et al., 2022; Zhang et al., 2012; Zrachya et al., 2007), (ii) it is also required for viral movement and spreading of the virus through the plant (Moshe et al., 2015; Padidam et al., 1996; Rojas et al., 2001; Rothenstein et al., 2007) and (iii) it is involved in the regulation of other host defence responses different than RNA silencing (Bar-Ziv *et al.*, 2012; Roshan et al., 2018).

V2 from *Beet curly top virus* (BCTV), the model species for the *Curtovirus* genus, suppresses PTGS by disrupting the RDR6/SGS3 pathway and induce a hypersensitive response-like (HR-like) when expressed from *Potato virus X* (PVX) vector. The protein presents two hydrophobic domains and a putative phosphorylation motif, which are essential for systemic infection, silencing suppression activity, and nuclear localization of the protein (Luna et al., 2017; Luna et al., 2020).

*Beet curly top Iran virus* (BCTIV) is one of the most important causal agents of beet curly top disease (BCTD) in Iran, and it has been recently described as the type species of the genus *Becurtovirus* (Heydarnejad et al., 2007; 2013). Its genome presents a unique organization: similarly to *Mastrevirus*, it contains two intergenic regions, one large (LIR) and another small (SIR), located at opposite sides of the genome, but BCTIV LIR includes a sequence capable of forming a stem-loop structure and a novel nonanucleotide (TAAGATT/CC) with a unique nick site. The BCTIV genome comprises three virion-sense (V1, V2, and V3) and two complementary-sense (C1 and C2) ORFs. Based on a comparison of nucleotide sequence identity of individual genes, the three virion-sense ORFs V3, V2 and V1 are similar to their positional homologs in the Curtoviruses, whereas C1 and C2 are more similar to the ones from Mastreviruses (Bozorgi et al., 2017; Yazdi et al., 2008) (Fig. 1A).

**Fig.1:**
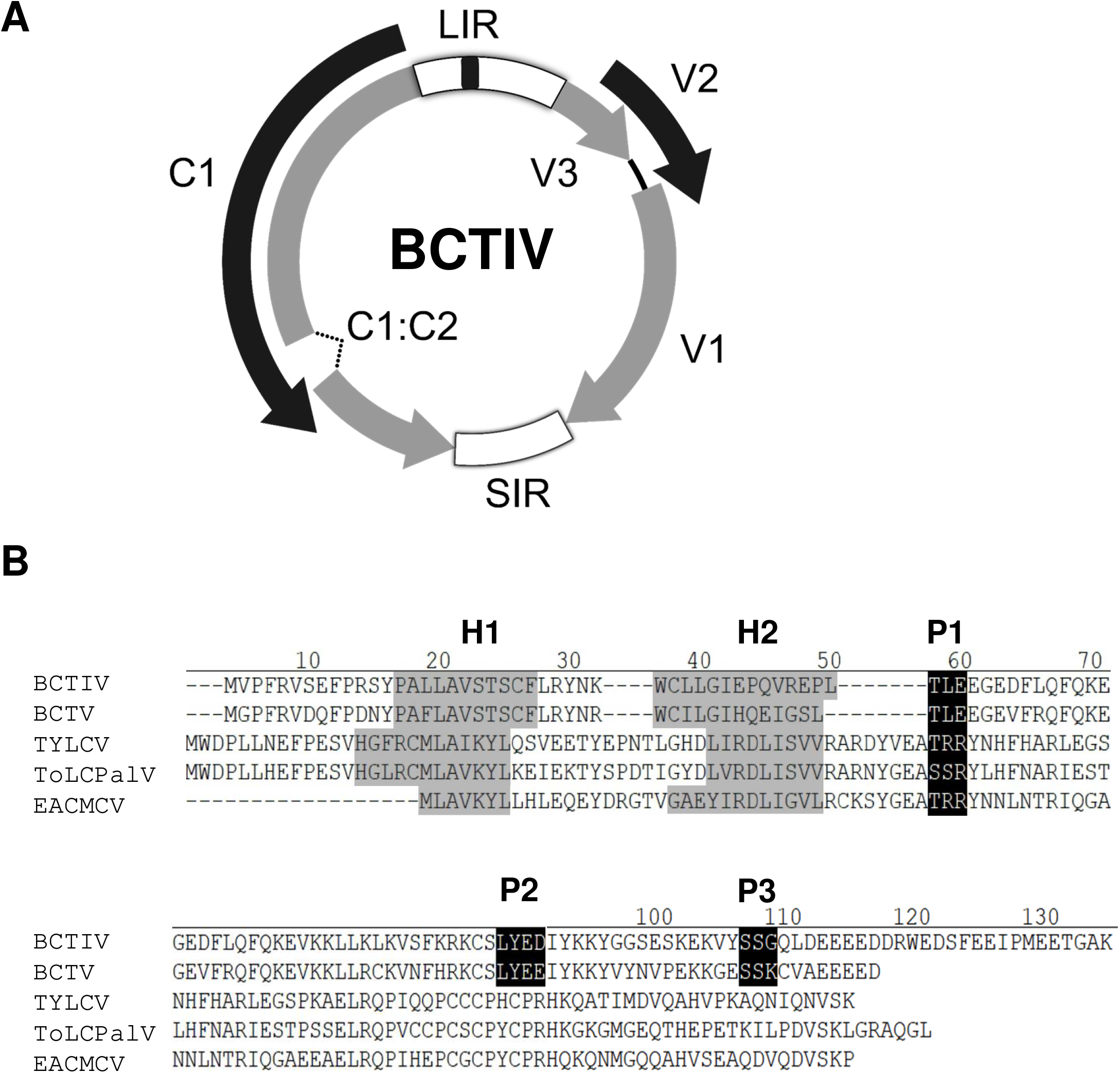
(A): *Beet curly top Iran virus* (BCTIV) genome structure. Arrows represent open reading frames (ORFs). LIR: long intergenic region; SIR: short intergenic region. Common region inside LIR is depicted in black (B): Alignment of the amino acid sequences of the V2 proteins from the becurtovirus beet curly top Iran virus (BCTIV; AFK14083), the curtovirus beet curly top virus (BCTV; M24597) and the begomoviruses tomato yellow leaf curl virus (TYLCV; X15656), tomato leaf curl Palampur virus (ToLCPalV; CAP03292) and East African cassava mosaic Cameron virus (EACMCV; AF112354). Gaps (-) were introduced to optimize the alignment. The positions of the predicted putative phosphorylation motifs P1 (protein kinase CK2/protein kinase C), P2 (protein kinase CK2) and P3 (protein kinase C) are depicted in white letters inside black boxes. The hydrophobic domains (H1 and H2) are shadowed in gray.

This work aims to determine the role of BCTIV V2 in pathogenicity and its function as an RNA silencing suppressor. This knowledge will help to study further the molecular mechanisms involved in the infection process and develop strategies for decreasing the impact of this disease in agriculture.

## 2. Materials and methods

### 2.1. Microorganisms, plant material, and growth conditions

Manipulation of *Escherichia coli* strains were carried out according to standard methods (Sambrook, 2001). *E. coli* strain DH5-α was used for subcloning. The *Agrobacterium tumefaciens* GV3101 strain was used for the agroinfiltration and agroinoculation/infection assays.

Plants used in this study were wild-type *Nicotiana benthamiana* and 16c *N. benthamiana* line (transgenic plants constitutively expressing green fluorescent protein [GFP] (Ruiz et al., 1998). Plants were grown in chambers at 24°C in long-day conditions (16 h light/8 h dark) before and after agroinfiltration/infection.

### 2.2. Bioinformatic analysis

The ClustalW algorithm was used to align V2 and V3 homolog proteins. The prediction of the nuclear localization signals was made by NLS Mapper online tool (https://nls-mapper.iab.keio.ac.ip/cgi-bin/NLS_Mapper_form.cgi).

### 2.3. Plasmids and cloning

The sequences of the primers used in this study are listed in Table S1. All PCR-amplified fragments cloned in this work were fully sequenced.

#### Generation of binary vectors

Gateway-compatible oligonucleotide primers BIV2Fw and BIV2RvSt or BIV2RvNoSt were used to amplify BCTIV V2 full-length ORF, with or without stop codon, respectively, and to generate entry clones by performing BP recombination reactions between the pDONR™ vector (Gateway™ pDONR™/Zeo Vector) and the attB PCR products. Subsequently, these entry clones were used for LR recombination reactions with Gateway™ destination binary vectors: pGWB2 yielding pGWB2-IVV2, for their expression in plants from a 35S promoter, and pGWB5 and pGWB6 yielding pGWB5-IVV2 and pGWB6-IVV2, to express V2 fused to GFP in the carboxyl- or the amino terminal region of the viral protein respectively (Nakagawa et al., 2009).

#### Generation of PVX expression vectors

To generate the PVX expression vector, BCTIV V2 complete ORF was amplified using primers ClaI V2 F and SalI V2 R, and subcloned into the pGEMT-easy vector (Promega Corp., Madison, WI, U.S.A.) to yield pGEMT-IVV2. Restriction fragments *ClaI-SalI* from this plasmid containing the complete ORF were cloned into PGR107 (Jones et al., 1999), yielding the corresponding PVX expression vector PVX-IVV2.

#### Generation of V2 mutant BCTIV virus

As a first step, a 1.3mer genome-length infective BCTIV clone was obtained from the previously constructed BCTIV dimer infective clone (accession number: JQ707949)(Heydarnejad et al., 2013). The dimer was first digested by *KpnI* (TaKaRa, Kyoto, Japan) to release a monomer, and subsequently with *Eco*RI and *EcoRV* (TaKaRa, Kyoto, Japan) to release a 0.3mer fragment. This 0.3mer fragment was cloned into *KpnI/EcoRI* sites of pGreen0229 binary vector (Hellens et al., 2000), yielding pGreen 0.3 BCTIV. Finally, the monomer obtained by *Kpn*I digestion was cloned into *Kpn*I site of pGreen 0.3 BCTIV, to obtain the 1.3mer genome-length infectious clone pGreen 1.3 BCTIV.

BCTIVV2 mutant virus was then generated by site-directed mutagenesis using Nyztech Site-Directed Mutagenesis kit (Nyztech, Lisboa, Portugal) (Liu & Naismith, 2008) with specific primers IVV2 stop F and IVV2 stop R (Table S1) designed to introduce a stop codon in the V2 gene.

### 2.4. Agroinfiltration and infection assays

For local and systemic PTGS suppression assays and PVX infection, *N. benthamiana* leaves were agroinfiltrated as previously described (Luna et al., 2012) Long-wave UV lamp was used to detect GFP fluorescence (Black Ray model B 100 AP, Upland, United States). Pictures were taken using EOS 5D Canon digital camera (Tokyo, Japan)

For BCTIV infection, *N. benthamiana* plants at two leaves stage were agroinoculated as described by (Elmer et al., 1988) Elmer et al., 1988 (final OD600 =0.2).

### 2.5. Subcellular localization

For subcellular localization, *A. tumefaciens* was transformed with binary vectors containing BCTIV V2 or BCTV V2 fused to GFP in the carboxyl- or the amino terminal region of the viral protein. *N. benthamiana* leaves were agroinfiltrated with cultures at OD600 0.5–1. Fluorescence was detected in epidermal cells 36 hours after infiltration using a confocal microscope (Zeiss LSM 880).

### 2.6 Analysis of nucleic acids and proteins

Nucleic acid manipulation was performed according to standard methods(Sambrook, 2001).

For BCTIV replication and infection analyses, plant DNA was extracted from the infiltrated (local) or the apical (systemic) leaves of the infected plants at 6 or 28 days post-infiltration (dpi), respectively, digested with *DpnI* to remove bacterial DNA in the infiltrated tissues (local infection) and then subjected to qPCR analysis using primers BCTIV RTF and BCTIV RTR primers (Table S1), and *N. benthamiana Elongation Factor α* (EF1 α) as normalizer (Rotenberg et al., 2006).

Expression of BCTIV V2 and BCTV V2 in agroinfiltrated tissues was determined by semiquantitative RT-PCR using the specific primers *Cla*I V2 F and *Sal*I V2 (Table S1). EF1 α was used as an internal gene control.

For western blot analysis, 120 mg of leaf tissue per sample were used. Total protein was extracted by 2X Laemmli buffer and separated using sodium dodecyl sulfate (SDS)–10% polyacrylamide gel electrophoresis (PAGE), then transfered to PVDF membrane (Immobilon-P, Millipore, MA, United States) by a semi-dry electrotransfer (Bio-Rad, Hercules, California, USA) [36]. It was then probed by an anti-GFP mouse monoclonal antibody (1:600, clone B-2; sc-9996, Santa Cruz Biotechnology, Dallas, Texas, USA). Anti-mouse IgG conjugated by peroxidase (A9044, Sigma-Aldrich, Missouri, USA) was used as a secondary antibody at 1:80,000 (V/V) dilution.

## 3. Results

### 3.1. V2 from BCTIV is a strong local PTGS suppressor

Amino acidic comparison among V2 from BCTIV with the homolog proteins from curtoviruses and begomoviruses revealed that BCTIV V2 contains a conserved CK2/PKC (protein kinase CK2/protein kinase C) phosphorylation motif (hereafter named P1) present in all V2 proteins and two putative CK2 and PKC phosphorylation motifs predicted in BCTV V2 (named P2 and P3, respectively) are also present in BCTIV V2, but not in begomoviral proteins (Fig.1B). Besides those phosphorylation motives, all V2 proteins have similar hydrophobic profiles displaying two hydrophobic regions at the N-terminus (hereafter named H1 and H2), followed by a long hydrophilic region at the C-terminus (Fig. 1B). These hydrophobic regions, along with P1, but not P2 or P3, have been proved to be essential for PTGS suppression activity and pathogenicity in begomovirus and curtovirus (Chowda-Reddy et al., 2008; Luna et al., 2020). Besides those conserved regions, the bioinformatic analysis identified a nuclear localization signal (NLS) present in BCTIV V2 as well as in V2 from the curtovirus BCTV protein sequences while it was absent in most of the begomoviral ones. (Supp. Fig. 1)(Dingwall & Laskey, 1991; Robbins et al., 1991).

To test if BCTIV V2 functions as a PTGS suppressor, we carried out transient expression assays in *N. benthamiana* plants. Leaves were co-infiltrated with *Agrobacterium* cultures expressing GFP (35S:GFP) and BCTIV V2 (thereafter named IVV2). As controls, leaves were co-infiltrated with 35S:GFP and the empty vector (EV) (negative control) or a plasmid expressing the viral silencing suppressor V2 from BCTV (thereafter named BCV2) (positive control). Leaves were observed under UV light at 6 dpi. GFP fluorescence was still strong in the tissues co-infiltrated with 35:GFP and IVV2, as in leaves infiltrated with BCV2. Conversely, in the patches infiltrated with the empty vector, fluorescence had disappeared almost completely (Fig. 2A). These differences in green fluorescence were confirmed by western blot analysis using an anti GFP antibody. As expected, GFP protein accumulation in tissues infiltrated with IVV2 at 6 dpi was considerably higher than in tissues co-infiltrated with GFP and the empty vector. This enhancement of GFP expression was as strong as the one produced by BCV2, proving that BCTIV V2 is also a strong PTGS suppressor at a local level (Fig. 2B). Expression of viral genes in the infiltrated tissues was confirmed by semiquantitative RT-PCR (Fig. 2C).

**Fig. 2:**
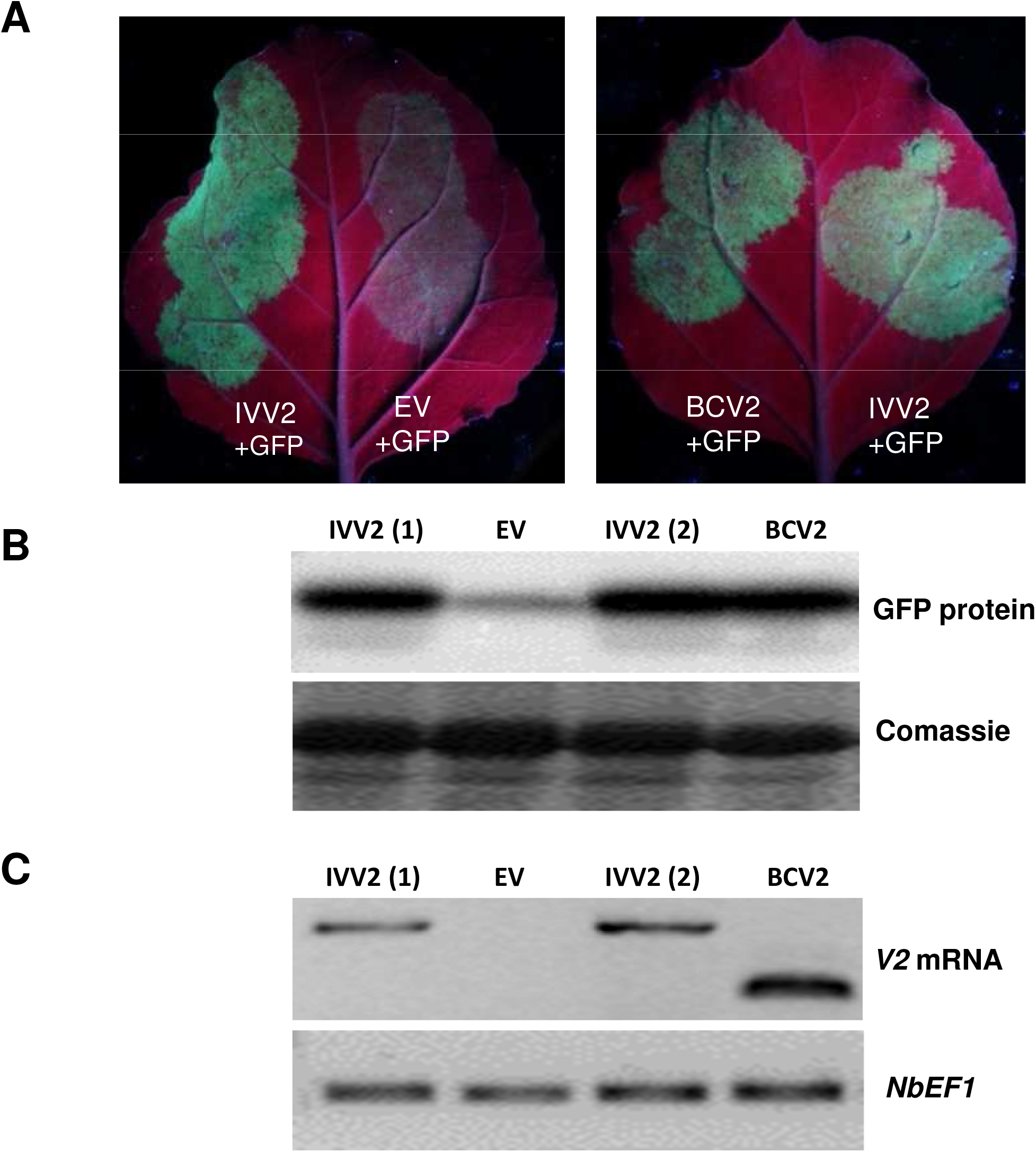
Local PTGS suppression assay in wild type *Nicotiana benthamiana*. (A) Leaves co-infiltrated by a mixture of two *Agrobacterium tumefaciens* cultures expressing GFP and V2 from beet curly Iran top virus (IVV2), under UV light at 6 days postinfiltration (p.i.). V2 from beet curly top virus (BCV2) and the empty vector (EV) were used as controls. (B) Western blot was performed by anti-GFP antibody. Coomassie blue staining of SDS-PAGE gel is shown as a loading control. Four to six plants were agroinfiltrated per experiment. Similar results were obtained in three independent experiments. (C) V2 viral protein transcription was confirmed by RT-PCR and *Elongation factor alpha (NbE1Fa)* was used as an internal control.

### 3.2. V2 from BCTIV causes a delay in the long-distance spread of RNA silencing

To determine the effect of BCTIV V2 on systemic PTGS, the constructs expressing GFP and IVV2 or BCV2 were agroinfiltrated in leaves of transgenic 16c *N. benthamiana* plants. Co-infiltration with 35S:GFP and P19 or the empty vector were used as positive or negative controls, respectively.

Fluorescence was monitored under UV light to detect the initiation of systemic silencing in newly emerging leaves, from 4 to 30 dpi. To evaluate the silencing progress, we applied the arbitrary silencing index described in Luna et al., 2012 (Supp. Fig. 2). Systemic silencing in emerging upper leaves was not observed until 8 dpi. After 18 dpi, systemic GFP silencing was observed in plants co-infiltrated with 35S:GFP and the empty vector and in plants infiltrated either with IVV2 or BCV2, suggesting neither of these proteins is able to suppress the spreading of the GFP silencing. As expected, in plants infiltrated with P19, no systemic silencing was observed. However, there was a clear reduction in the extension of the silenced tissues in the IVV2 and BCV2 infiltrated plants compared to ones infiltrated with the empty vector, indicating that both proteins, IVV2 and BCV2 can produce a slight delay in the silencing spreading, although none of them seems to be blocking it completely. This delay was maintained until 30 dpi (Supp. Fig. 2, Fig. 3A).

**Fig. 3:**
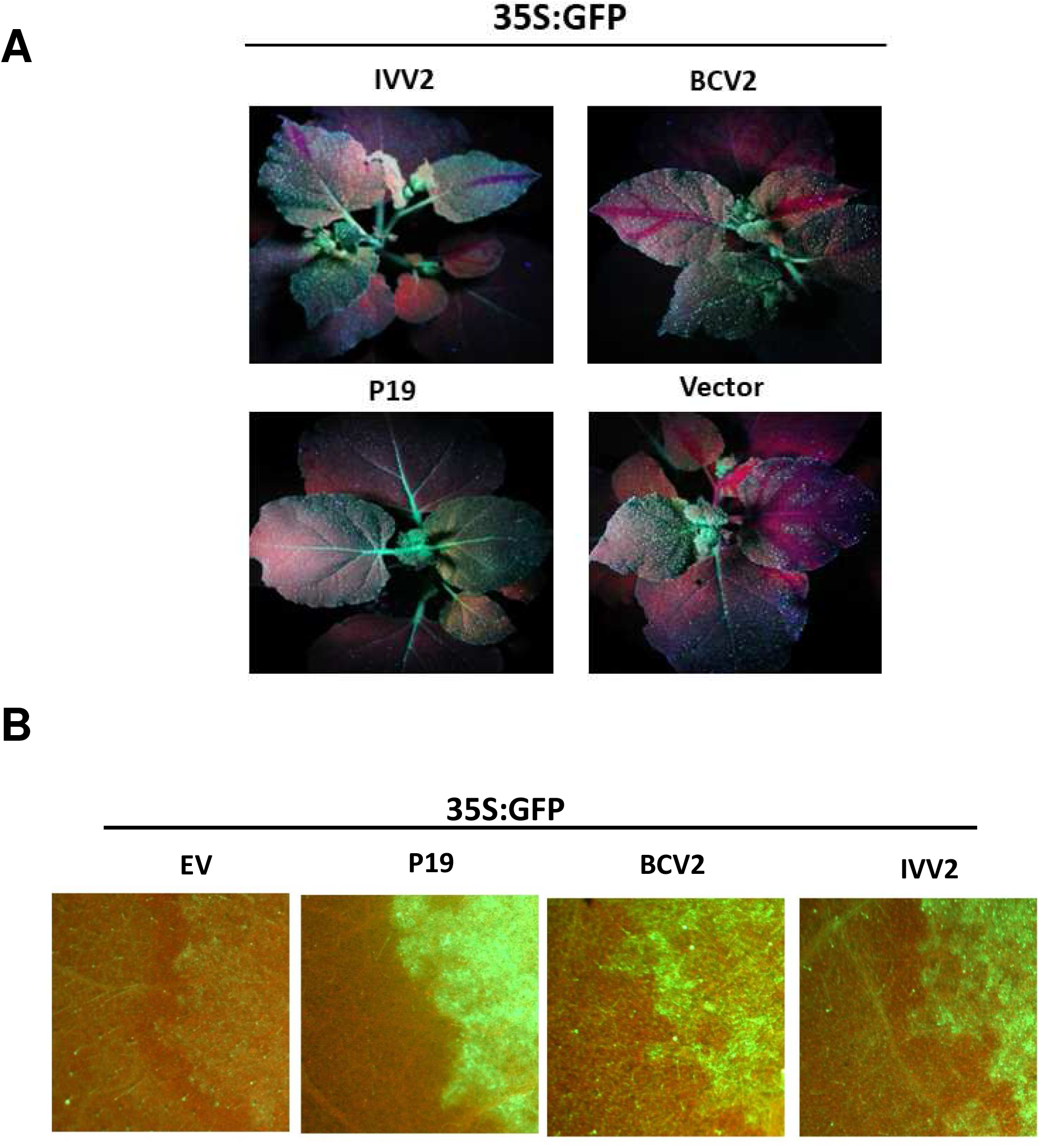
Effect of V2 from beet curly top Iran virus on long distance and short range spread of RNA silencing in 16c *Nicotiana benthamiana* plants. (A) 16c plants were agroinfiltrated with two *A. tumefaciens* cultures expressing GFP and V2 from beet curly top Iran virus (IVV2). V2 from beet curly top virus (BCV2), P19 and the empty vector (EV) were used as controls. Agroinfiltrated16 c plants were observed under UV light at 18 days post inoculation (dpi). (B) GFP expression in the cells surrounding the agroinfiltrated area at 6 dpi. Four to six plants were agroinfiltrated per experiment. Similar results were obtained in three independent experiments.

### 3.3. V2 from BCTIV cannot suppress short-range (cell-to-cell) spread of gene silencing

Agroinfiltrated 16c plants were also employed to study the effect of BCTIV V2 on the short-range movement of the RNA silencing. For that, GFP expression was monitored with UV light in the cells surrounding the infiltrated tissues at 6 dpi, to check if the silencing signal triggered by the GFP overexpression had been able to exit from the infiltrated area and cause the appearance of a red ring around the patch (Himber et al., 2003). In the plants co-infiltrated with 35S:GFP and the empty vector, the red ring was observed at 6 dpi (Fig. 3B). This red ring was also present around the patches infiltrated with BCV2, but not in those infiltrated with P19, as previously described (Himber et al., 2003; Luna et al., 2017; Silhavy et al., 2002). In the plants infiltrated with IVV2 a red ring was also visible at 6 dpi, showing that this viral protein, as its Curtoviral counterpart, cannot block the cell-to-cell movement of the silencing signal to adjacent cells.

### 3.4. V2 from BCTIV triggers HR-like response when expressed from a PVX derived vector

As mentioned above, ectopic expression of V2 proteins from *Begomovirus* and *Curtovirus* from a potato virus X (PVX)-derived vector causes localized cell death in the infiltrated tissues and produces systemic necrosis associated with a hypersensitive response-like (HR-like) phenotype in *N. benthamiana* (Luna et al., 2012; Luna et al., 2017; Mubin et al., 2010; Roshan et al., 2018; Sharma & Ikegami, 2010). To study the possible role of BCTIV V2 in pathogenicity, we used the PVX-derived vector pGR107, to express the viral protein in *N. benthamiana* plants (PVX-IVV2). Agroinfiltration with the empty PVX vector was used as a negative control, whereas infiltration with PVX expressing either BCTV-V2 (PVX-BCV2) or the nonstructural protein (NSs) from *Tomato spotted wilt virus* (PVX-NSs) were used as positive controls (Bucher et al., 2003; Luna et al., 2017; Takeda et al., 2002).

At 5 dpi, tissues infiltrated with the empty vector developed a local yellowing, typical for PVX, while tissues infiltrated with PVX-IVV2 showed intense local necrosis, as the infiltrated with the positive controls: PVX-NSs and PVX-BCV2 (Fig.4A). At 16 dpi, plants inoculated with PVX expressing any of the viral proteins, including IVV2, showed severe systemic necrosis and posteriorly did not recover from a viral infection, while the ones inoculated with the empty PVX vector were almost asymptomatic. Interestingly, although the three viral proteins induce necrosis, the intensity and spreading of this necrosis is clearly larger for PVX-NSs and PVX-IVV2 (Fig. 4B).

**Fig. 4:**
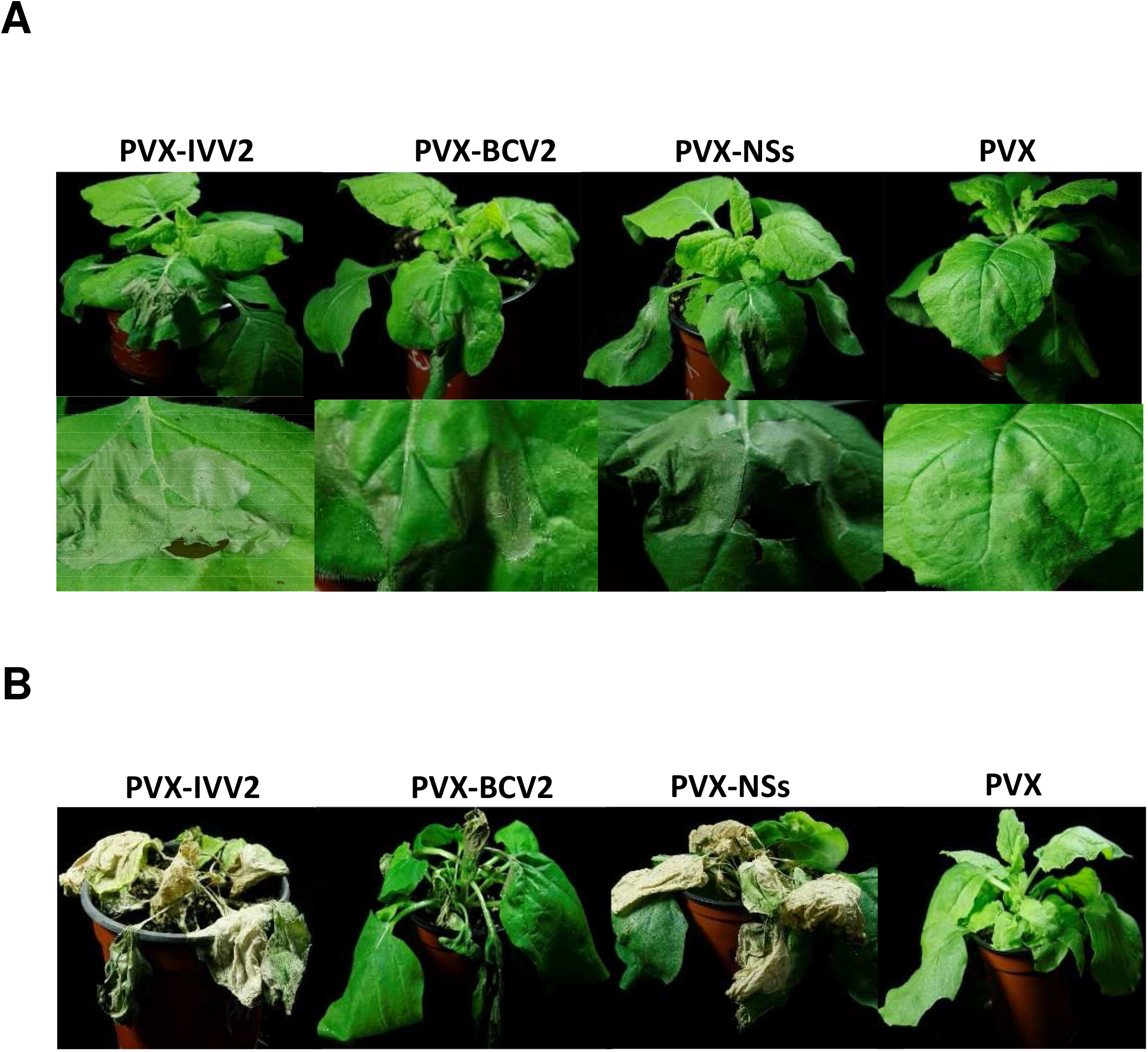
Local and systemic symptoms induced by potato virus X (PVX) expression of V2 from beet curly top Iran virus (PVX-IVV2) in *Nicotiana benthamiana* plants. Empty PVX vector (PVX) was used as negative control. Recombinant PVX viruses expressing either the nonstructural protein (NSs) from tomato spotted wilt virus (PVX-NSs) or the V2 protein from beet curly top virus (PVX-BCV2) were used as positive controls. (A) Local symptoms induced in the infiltrated area at 6 dpi. (B) Representative plants showing systemic symptoms at 16 dpi. Four to six plants were agroinfiltrated per experiment. Similar results were obtained in three independent experiments.

### 3.5. V2 from BCTIV localizes in the nucleus and perinuclear region

To obtain additional information on the function of BCTIV V2, we examined the subcellular localization of the protein, expressing GFP-fused versions of BCTIV V2 (both amino- and carboxyl-terminal fusions: GFP-IVV2 and IVV2-GFP respectively) in *N. benthamiana* leaves. BCTV V2 fused to GFP in the amino-terminal region of the viral protein (GFP-BCV2), and free GFP (GFP) were used as controls. Thirty-six hours after the agroinfiltration, tissues were collected and visualized using a confocal microscope. BCTIV V2 seems to have a similar subcellular localization to other V2 proteins, including its curtoviral counterpart (Guo et al., 2015; Kotlizky et al., 2000; Luna et al., 2017; Luna et al., 2017; Takeda et al., 2002; Wang, et al., 2019; Zhan et al., 2018). Accumulation of the protein is detected in the nucleoplasm and in the cell periphery, where it forms punctate fluorescent bodies. A closer examination shows that V2 accumulates in a discrete nuclear structure that could correspond to the Cajal body (Fig.5, Supp. Fig 3).

**Fig. 5:**
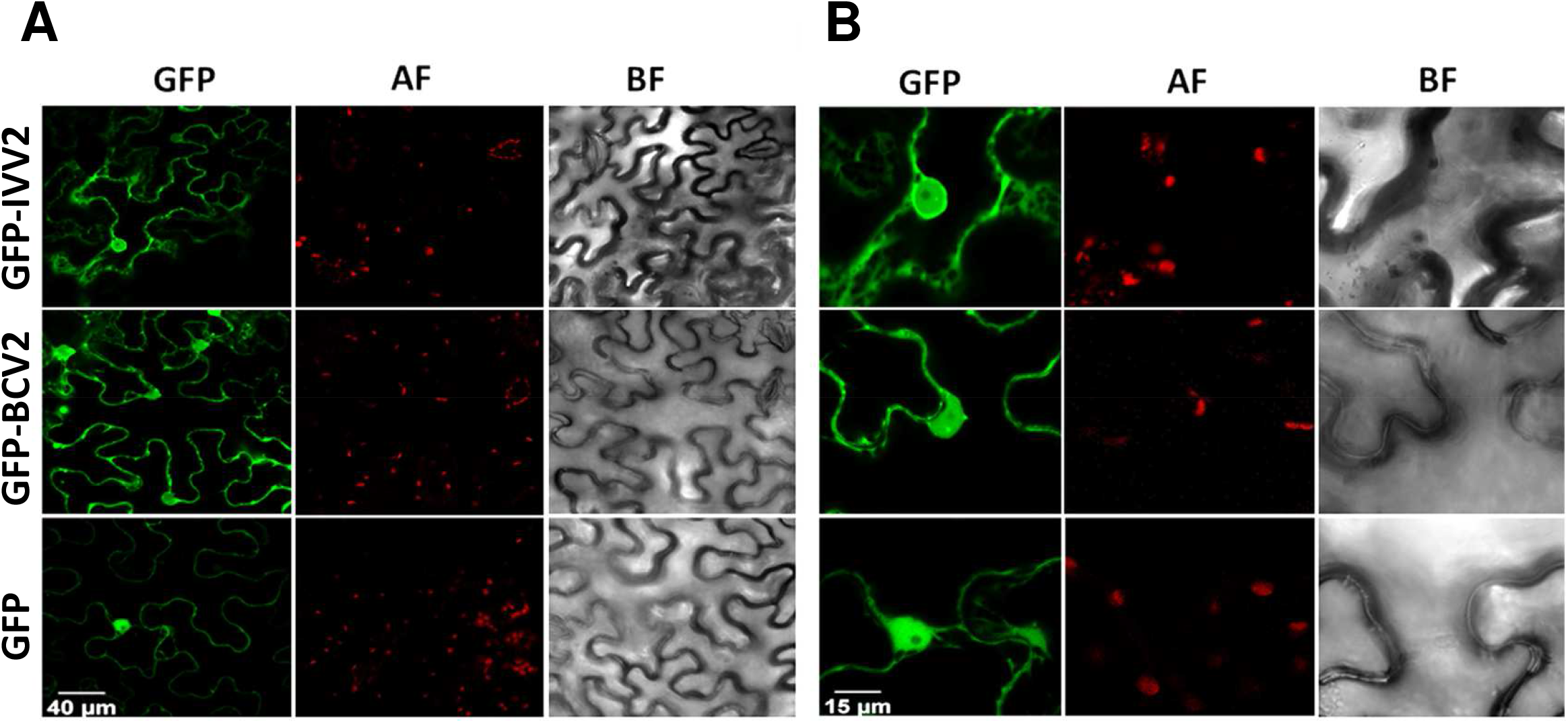
Subcellular localization of V2 from beet curly top Iran virus fused to GFP in epidermal cells of *Nicotiana benthamiana*. (A) Leaves were agroinfiltrated with a construct expressing the 35S:GFP (GFP), 35S:GFP-V2 fusion protein from beet curly top Iran virus (GFP-IVV2) or the 35S:GFP-V2 fusion protein from beet curly top virus (GFP-BCV2). Samples were observed under the confocal microscope at 36 hours post infection. (B) Close up confocal images of the observed areas. GFP fluorescence (GFP), autofluorescence (AF) and the bright field channel (BF) are shown

### 3.6. V2 from BCTIV is essential for systemic, but not for local infection of BCTIV

To complete the characterization of BCTIV V2, we determined its relevance for the systemic viral infection. To achieve this, we generated a BCTIV infective clone containing a stop mutation in the eighth codon of V2 ORF. This mutation also produced a change in the overlapping V3 gene, replacing a non-conserved arginine in the 33th position to leucine. *N. benthamiana* plants were agroinoculated with the V2 mutant BCTIV (BCTIV-V2stp) or with the wild-type BCTIV (wt BCTV) as a positive control. Agroinoculation with the empty vector was used as a negative control. Symptoms were detectable in plants infected with wt BCTIV at 10 dpi, while plants infected with the V2 mutant remained asymptomatic. At 28 dpi wt BCTIV-infected plants showed the typical BCTIV symptoms (stunting, leaf curling, vein swelling, etc.), whereas those agroinoculated with BCTIV-V2stp did not develop any symptoms (Fig. 6A).

**Fig. 6:**
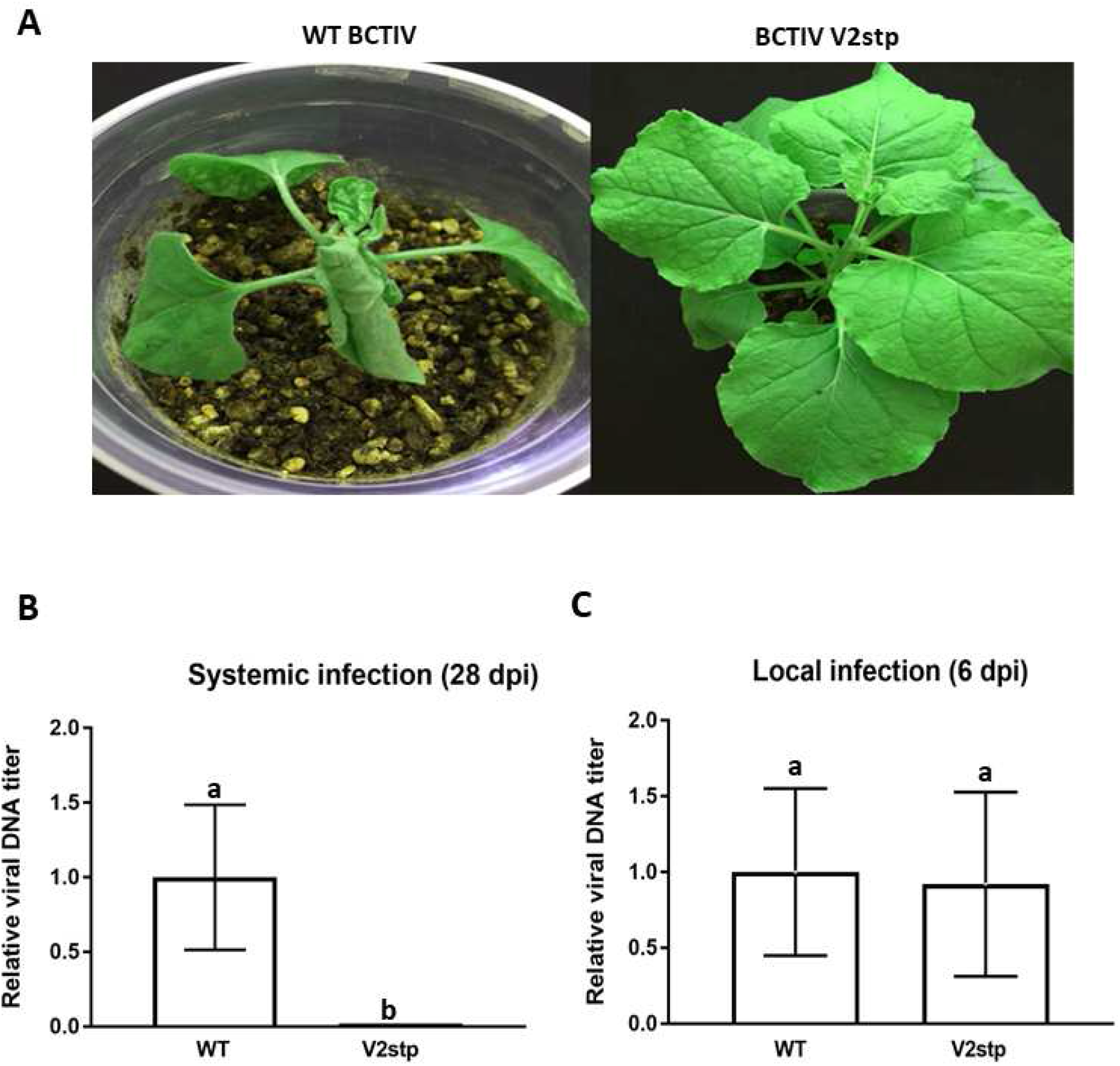
Infection of *Nicotiana benthamiana* plants with wild type beet curly top Iran virus and the V2 stop mutant virus. Plants were agroinoculated with wild-type (wt BCTIV) or V2 mutant BCTIV (BCTIV V2stp) infective clones. (A) Representative plants showing BCTIV symptoms at 28 dpi. Relative viral DNA accumulation in apical leaves (systemic infection) at 28 dpi (B) or in the infiltrated leaves (local infection) at 6 dpi (C). BCTIV accumulation was measured by qPCR after *Dpn*I treatment to remove bacterial DNA (6dpi). DNA levels were normalized to *Elongation factor alpha* (NbE1Fa) and are presented as the relative amount of virus compared with the amount found in wild-type BCTIV samples (set to 1). Bars represent mean values ± standard error (SE) for 6 pools of two leaves. Mean values marked with different letter (a or b) indicate results significantly different from each other, as established by Student’s t-test α <0.001) virus titer.

To determine if the absence of symptoms correlated with an absence of viral DNA, total DNA was extracted from the newly emerging leaves of the inoculated plants at 28 dpi, and viral DNA was quantified by quantitative real-time PCR (qPCR). No BCTIV DNA was detected in the apical leaves of plants inoculated with BCTIV V2 stop (Fig. 6B), indicating that BCTIV V2 stop could not infect plants systemically. To test if V2 is required for viral replication, we performed a local infection assay. Total DNA was extracted at 6 dpi from the infiltrated leaves and viral DNA was quantified by qPCR. No differences in the amount of viral-DNA accumulated in the infected tissues were detected in the leaves infected by wt BCTIV or BCTIV-V2stp (Fig. 6C). This result suggested that BCTIV V2 is required for viral movement but not for replication.

## 4. Discussion

BCTD is one of Iran’s most damaging sugar beet diseases caused by *Turncortovirus, Curtovirus*, and *Becurtovirus* members. These viruses share hosts, transmission mode, and vector, and as a result, the occurrence of mixed infections in the field is very frequent (Astaraki et al., 2020). BCTIV is one of the main causal agents. This *Becurtovirus* is a natural recombinant virus from *Curtovirus* and *Mastrevirus* ancestors. Up to date, no studies have been carried out to study the function of any of the ORFs identified in the viral genome (Heydarnejad et al., 2013; Yazdi et al., 2008). In order to better know the virus-plant interactions and the mechanisms of viral infection for this virus, we have characterized one of these ORFs, named V2. This gene plays an important role in viral infection and pathogenicity for many Begomoviruses and some Curtoviruses.

Given the conservation of protein domains/regions (P1, H1, or H2) present in all geminiviral V2 and involved in PTGS suppression and pathogenicity (Chowda-Reddy et al., 2008; Luna et al., 2020), it was expected that BCTIV V2 would have similar activities. The results obtained in local and systemic silencing assays (Figs 2 and 3) have proved that BCTIV V2: (i) is a strong PTGS suppressor at a local level, (ii) it produces a delay in spreading of the systemic silencing, but (iii) cannot avoid the cell-to-cell movement of the silencing signal to neighboring cells. Furthermore, considering its high sequence homology with BCTV V2 protein (63% identity), it is possible that, as the curtovirual protein, BCTIV V2 is suppressing PTGS by impairing RDR6/SGS3 silencing pathway (Luna et al., 2017).

As BCTV V2, BCTIV V2 expression in a PVX heterologous system caused the induction of necrotic lesions in *N. benthamiana* like the ones produced during the HR, both at local and systemic levels in the *N. benthamiana* plants inoculated. (Fig. 4A, B). These results suggest that both proteins are avirulence factors that are directly or indirectly recognized by a resistance gene to trigger a defense response based on HR, known as effector-triggered immunity (ETI) (Ferreira et al., 2021). It is remarkable that ectopic expression of BCTIV V2 from a 35S promoter did not cause any necrotic lesion in the infiltrated area in *N. benthamiana* leaves (similar results are observed when expressing BCTV V2) (Luna et al., 2017). This difference with the effect from a PVX vector, could be due to an increased expression of the viral protein in this heterologous system or maybe, as it has been suggested by some experiments performed by Aguilar and colleagues, a combination of the actions of the geminiviral suppressor being over expressed and the one already contained in the PVX genome, P25 (Aguilar et al., 2015). The differential participation of the functional domains of the BCTIV V2 protein in the HR-like response and the suppression of PTGS would need further study.

Depending on the silencing pathway steps that the viral suppressor is targeting, the protein needs to be in a specific localization inside the cell (Borges & Martienssen, 2015). According to our subcellular localization results, BCTIV V2 is localized in the nucleus and the perinuclear region, possibly associated with RE (Fig. 5A). This location has been reported for other V2 proteins that are silencing suppressors, from BCTV, tomato yellow leaf curl virus (TYLCV), apple geminivirus (AGV) and mulberry mosaic dwarf-associated virus (MMDaV) (Luna et al., 2017; Rojas et al., 2001; Yang et al., 2018; Zhan et al., 2018). Interestingly, in a closer examination of the nucleus, the GFP fused BCTIV V2 protein seems to be accumulating in sub-nuclear structures that resemble Cajal bodies (CBs) (Fig. 5B, supplementary figure 2). CBs are nuclear membrane-less bodies composed of proteins and RNA that play an important role in plant-virus interaction. It has been described that CBs are essential for systemic infection, are involved in viral systemic movement, and in the case of DNA viruses, are involved in the suppression of TGS and viral genome methylation (Ding & Lozano-Durán, 2020). Interestingly, V2 from the begomovirus TYLCV inhibits the methylation of the viral genome through its interaction with AGO4 in CBs (Chen et al., 2010; L. Wang et al., 2020; 2019). If *Becurtovirus* V2 also has a role in TGS suppression will require further analysis.

The results from the infection of *N. benthamiana* with a V2 BCTIV mutant showed that V2 is required for systemic infection. Although, the mutation also generated a change in a non-conserved aminoacid of V3, we cannot absolutely discharge the possibility that this change could impact in the ability of the virus to infect the plant (Supp. Fig. 4). The fact that no differences with the wild-type viruses are observed in local infection/viral replication pointed to a role of V2 in viral movement. Geminiviral V2 protein has been involved in the viral movement in other geminiviruses, such as *Curtovirus*, *Mastrevirus* and monopartite *Begomovirus* (Kotlizky et al., 2000; Padidam et al., 1996; Rojas et al., 2001; Zhai et al., 2022). The presence of NLS in BCTIV V2 and its subcellular localization suggest that V2 could be imported to the nucleoplasm and maybe participate, along with other viral proteins, in the viral movement outside the nucleus, as it has been proved for TYLCV V2 (Zhao et al., 2020).

Considering all the results, we conclude that BCTIV V2 is a functional homolog of curtoviral and begomoviral V2.

**Table 1:**
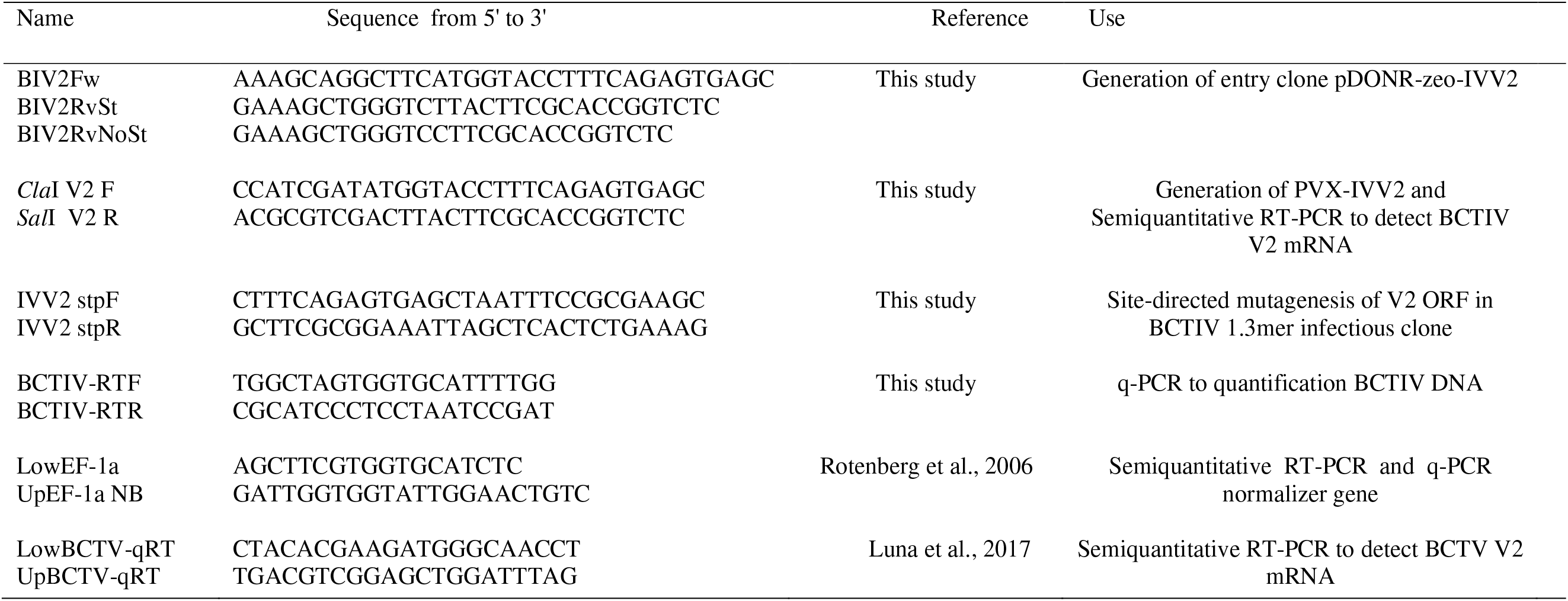
Primer used in this study.

## Declarations

### Ethical approval

This article does not contain any studies involving animals performed by any of the authors.

### Consent for publication

Not applicable

### Author contributions

Masoud Shams-bakhsh, Ana P. Luna, and Eduardo R. Bejarano designed and directed the project; Atiyeh Bahari, and Ana P. Luna performed the experiments, drafted the manuscript and designed the figures with support from Masoud Shams-bakhsh and Eduardo R. Bejarano. All authors discussed the results and contributed to the final manuscript.

## Acknowledgment

This researched was supported by Tarbiat Modares University, Iran and Universidad de Málaga, Spain.

## Availability of data and materials

The authors confirm that the data supporting the findings of this study are available within the article or its supplementary materials.

---

**Supplementary Fig. 1:**
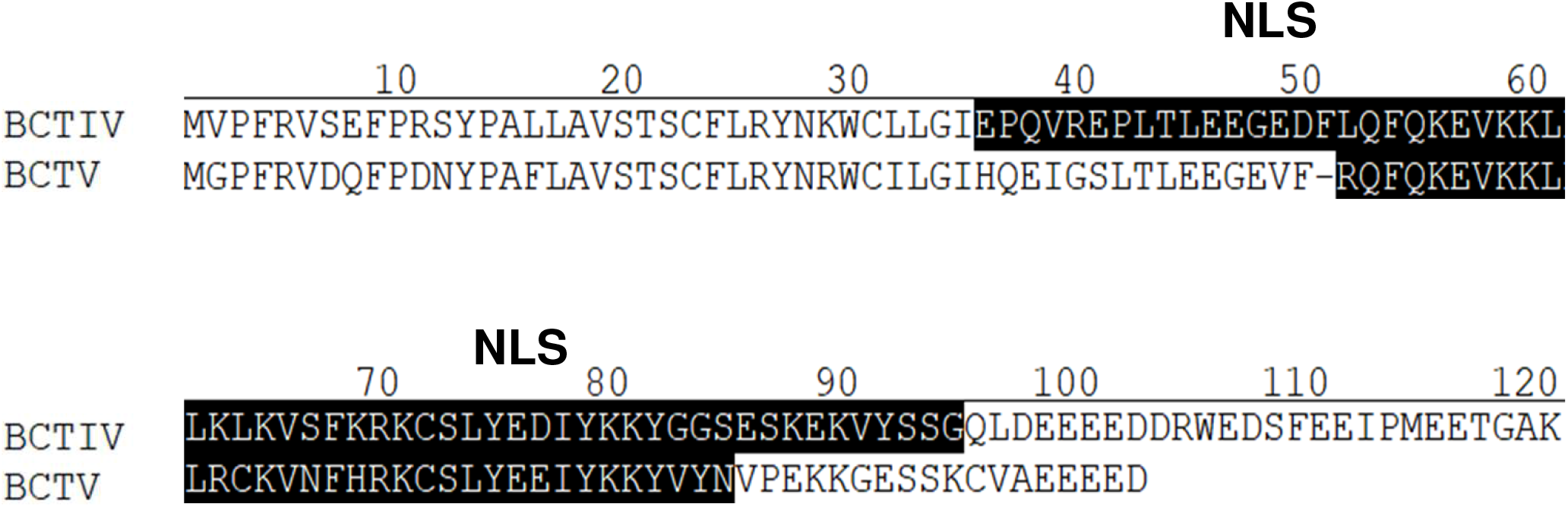
Alignment of the aminoacid sequences of the V2 proteins from the becurtovirus beet curly top Iran virus (BCTIV; AFK14083) and the curtovirus beet curly top virus (BCTV; M24597). Predicted Nuclear localization signal (NLS) are depicted in white letters inside black boxes.

**Supplementary Fig. 2:**
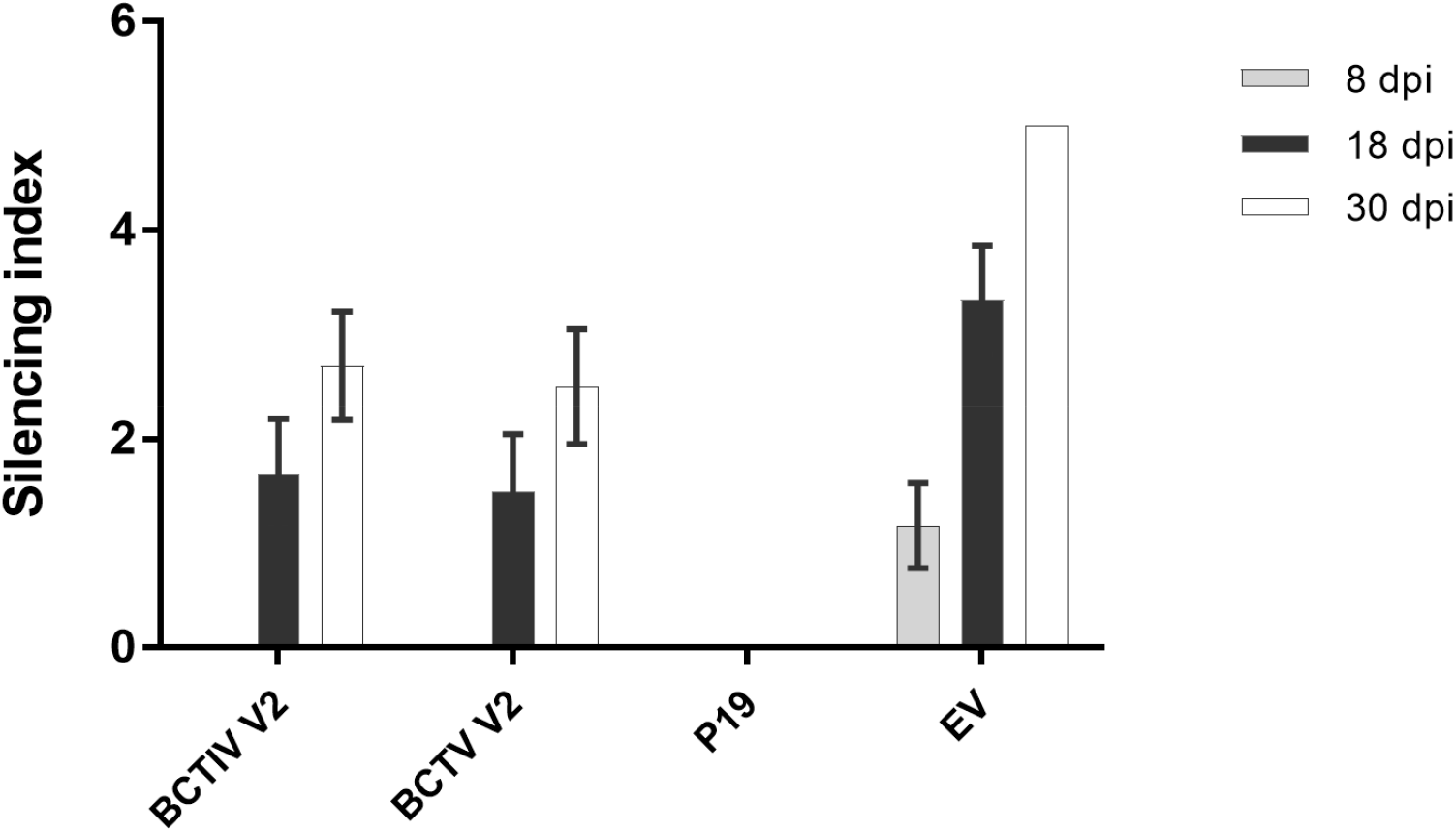
Levels of systemic silencing (Silencing index) of 16c *Nicotiana benthamiana* plants infiltrated with two different *Agrobacterium tumefaciens* cultures expressing GFP and the V2 proteins from beet curly top Iran virus (BCTIV) or beet curly top virus (BCTV V2) at 8, 18, and 30 days postinfiltration (dpi). The arbitrary silencing index described in Luna et al., 2012, was applied to visually categorize plants between six levels of systemic silencing, from 0 (plant with no silenced leaves) to 5 (plant with all of its leaves silenced). Values according to this index were assigned to individual plants, and the mean results per experiment on each observation day (6 plants per experiment) were represented graphically. Plants agroinfiltrated with constructs to express P19 or the empty vector (EV) were used as positive and negative controls respectively. Bars represent standard deviation

**Supplementary Fig. 3:**
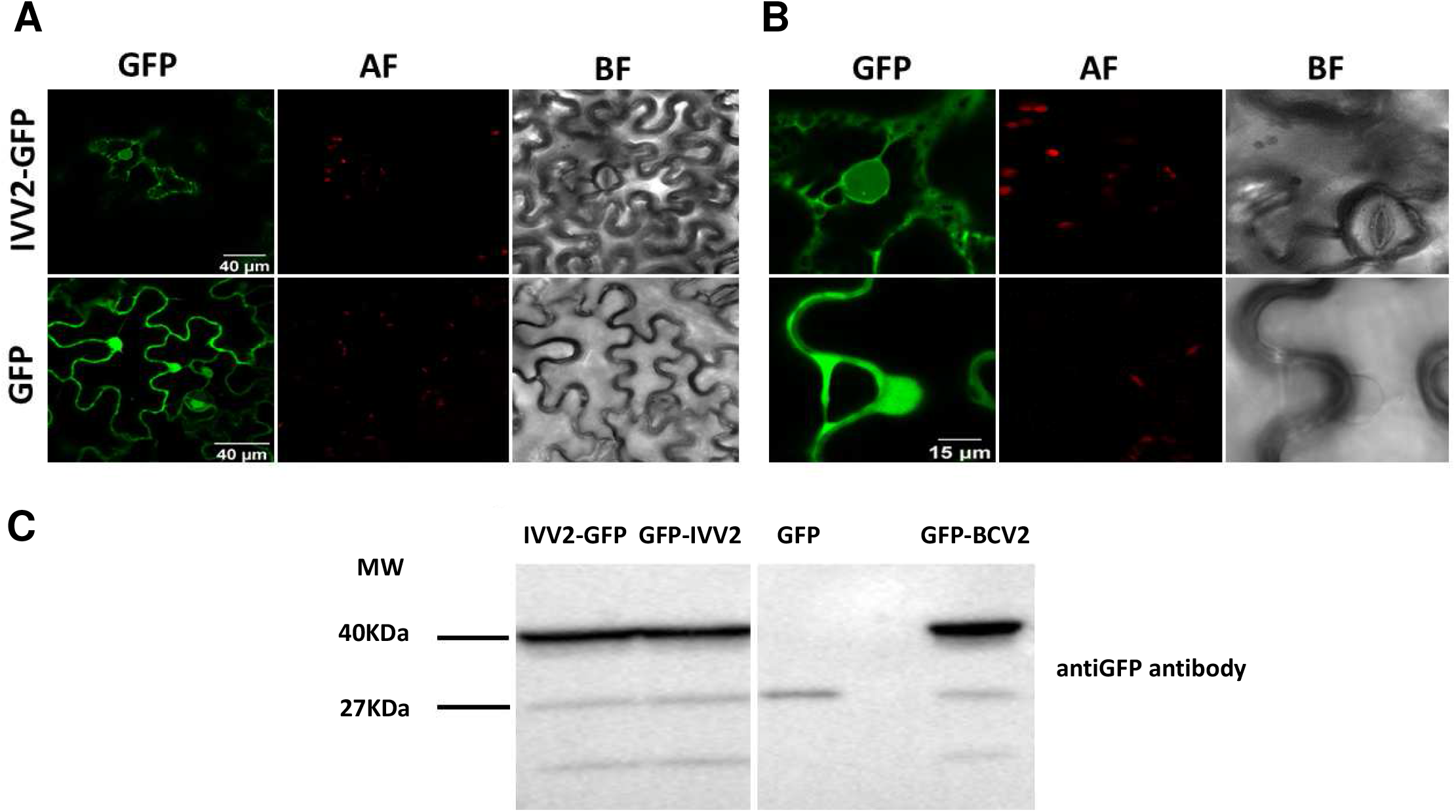
Subcellular localization of V2 from beet curly top Iran virus fused to GFP in epidermal cells of *Nicotiana benthamiana*. (A) Leaves were agroinfiltrated with a construct expressing the 35S:GFP (GFP), 35S:GFP-V2 or 35S:V2-GFP fusion proteins from beet curly top Iran virus (GFP-IVV2 or IVV2GFP respectively) or 35S:GFP-V2 fusion protein from beet curly top virus (GFP-BCV2). Samples were observed under the confocal microscope at 36 hours post infection. (B) Close up confocal images of the observed areas. GFP fluorescence (GFP), autofluorescence (AF) and the bright field channel (BF) are shown (C) western blot analysis of the observed with anti GFP antibody.

**Supplementary Fig. 4:**
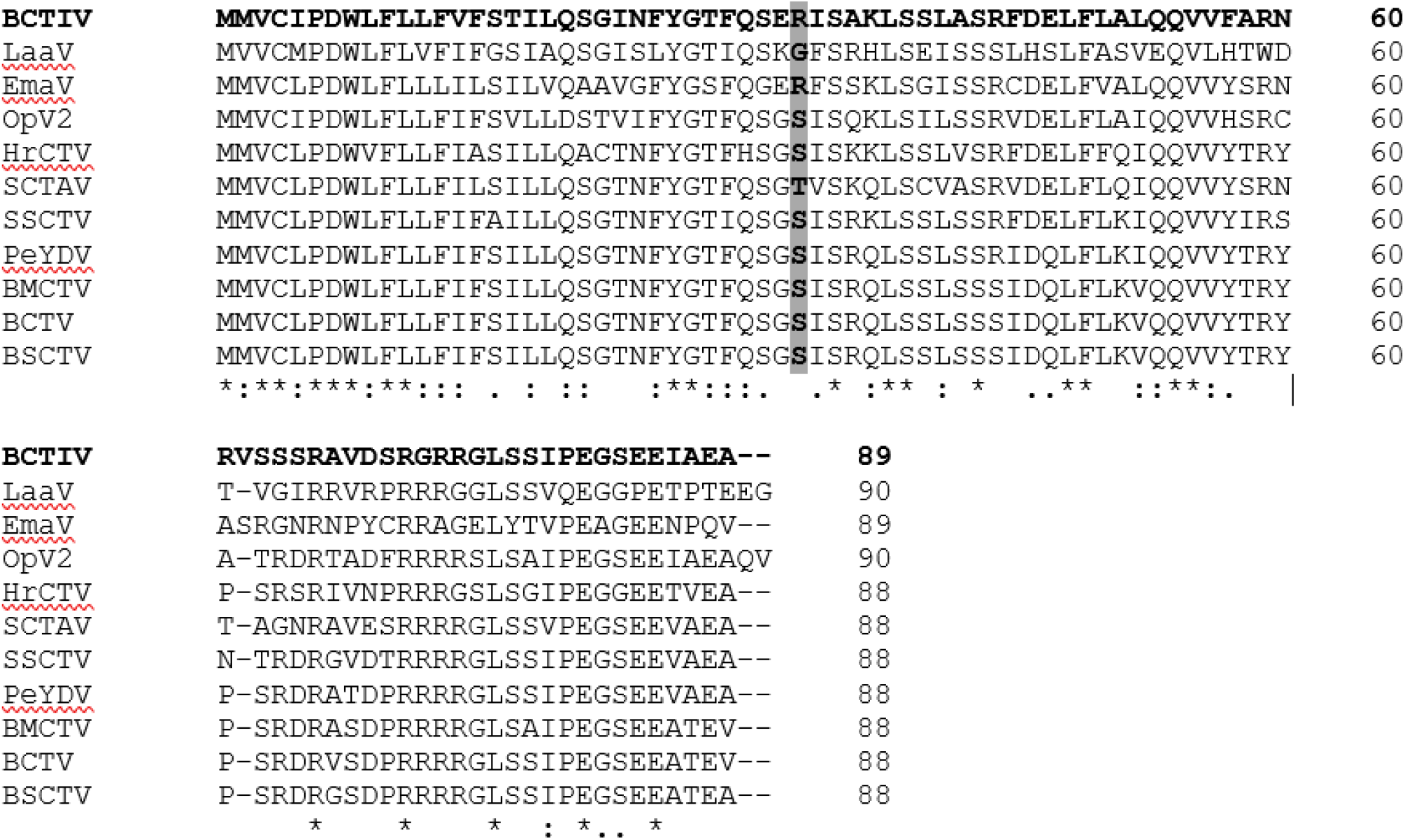
Alignment of the aminoacid sequences of the V3 proteins from the geminiviruses: beet curly top Iran virus (BCTIV; YP_001715617.1), Limeum africanum associated virus (LaaV; YP_009465967.1), Exomis microphylla associated virus (EmaV; YP_009465978.1), Opuntia virus 2 (OpV2; QTT61872.1), horseradish curly top virus (HrCTV; NP_066181.1), spinach curly top Arizona virus (SCTAV; YP_004207921.1), Spinach severe curly top virus (SSCTV; YP_003966133.1), pepper yellow dwarf virus-Mexico (PeYDV; YP_009362973.1), beet mild curly top virus (BMCTV; AAN06926.1), beet curly top virus (BCTV; NP_899663.1) and beet severe curly top virus (BSCTV; AAA20511.1). Changed aminoacid is highlighted in grey and bold letters.

